# Entomoscope 2.0 and ENIMAS 2.0: An Open-Source, AI-Integrated Platform for Rapid and Affordable Insect Digitization

**DOI:** 10.64898/2026.01.23.701290

**Authors:** Hossein Shirali, Nick Eric Böse, Markus Kramer, Jiahui Wang, Nathalie Klug, Ralf Mikut, Rudolf Meier, Christian Pylatiuk, Lorenz Wührl

## Abstract

The rapid decline of global biodiversity necessitates scalable and accessible tools for monitoring insect populations, yet the high cost and slow pace of specimen digitization remain significant bottlenecks. To address this challenge, we present the Entomoscope 2.0, an open-source platform that integrates a low-cost photomicroscope with an AI-integrated software suite, ENIMAS 2.0. The system combines optimized hardware with an end-to-end software module for digital specimen curation. This module comprises automated specimen cropping, background standardization, morphometric analysis using an Oriented Bounding Box (OBB) with a human-in-the-loop (HITL) supervision, and a flexible interface for rapid taxonomic screening using custom AI identification models. With a material cost of only 400 €, the system offers a cost-effective alternative to expensive commercial solutions. We compare results from the Entomoscope 2.0 with a high-end commercial system (Keyence) using 54 insect specimens and demonstrate the efficiency of the proposed AI workflow. Entomoscope 2.0 completed the whole digitization process in an average of 54.6 seconds per specimen, representing a 2.28-fold increase in speed over the Keyence system’s multi-step workflow (124.3 seconds). Crucially, all hardware specifications and construction manuals are freely available to support widespread adoption. By lowering financial barriers and accelerating research workflows, the Entomoscope 2.0 platform offers a practical solution to enable high-throughput digitization for researchers, educators, and citizen scientists.

## Introduction

The accelerating pace of global biodiversity loss demands scalable and accessible tools for monitoring insect populations. However, the resources for this task are limited: only a small fraction of insect biodiversity has been described, and the available expert capacity is insufficient to keep pace with urgent monitoring needs. Consequently, many groups remain critically understudied—the so-called ‘dark taxa’ (Hartop et al., 2022). The urgency of studying these taxa is compounded by the rapid reshaping of insect biodiversity driven by industrial agriculture and climate change (Raven and Wagner, 2021; Outhwaite et al., 2022). A primary bottleneck in addressing this knowledge gap is the acquisition of high-quality digital images required for taxonomic identification, morphological analysis, and biomass estimation. While powerful commercial photomicroscopy systems exist, their high cost is prohibitive for many research laboratories, educational institutions, and conservation initiatives, particularly in resource-limited regions where biodiversity is often greatest. Crucially, this high cost restricts most institutions to a single imaging unit, creating a bottleneck of serial processing. To tackle the massive scale of bulk samples, the field requires parallelization—the ability to operate multiple imaging stations simultaneously to multiply throughput—which is financially impossible with expensive commercial platforms. As highlighted by (Brydegaard et al., 2024), there is a critical need for “frugal methods” that empower researchers in the Global South to monitor biodiversity locally using affordable tools.

To address these challenges, entomology is rapidly moving towards integrated pipelines that combine robotics, imaging, high-throughput barcoding, and machine learning to tackle large-scale insect analysis (Wägele et al., 2022). This includes automated specimen handling (e.g., DiversityScanner or Biodiscover) (Ärje et al., 2020; Wührl et al., 2022), automated size-sorting (Ascenzi et al., 2025), high-resolution imaging (Klug et al., 2024), and AI-driven classification (Shirali et al., 2024; Caruso et al., 2025). Concurrently, the “reverse workflow” in molecular studies, in which specimens are first non-destructively barcoded (megabarcoding) (Srivathsan et al., 2021; Meier et al., 2025) and then morphologically validated (Hartop et al., 2024), further emphasizes the need for efficient, accessible, and non-destructive imaging tools. While genetic approaches provide taxonomic assignments, they do not capture phenotypical changes or age distributions. Furthermore, robust AI identification models for common species can significantly reduce the need for expensive DNA sequencing, but training these models requires massive, standardized image datasets.

While these high-throughput, fully automated systems are powerful, their inherent complexity and substantial cost restrict their widespread deployment, highlighting a persistent need for effective, semi-automated solutions that prioritize affordability and ease of use. The open-source hardware and Do-It-Yourself (DIY) movements present a promising pathway to democratize scientific instrumentation. Recent innovations include fully 3D-printed optical microscopes (Christopher et al., 2025a). While such solutions maximize affordability, the Entomoscope 2.0 distinguishes itself by utilizing a compound microscope design to ensure the high optical fidelity required for detailed taxonomy, while still prioritizing cost reduction.

Our previous work contributed to this effort by introducing the Entomoscope (Wührl et al., 2024). A collaboration between engineers, computer scientists, and entomologists then allowed for testing and improving the initial design, leading to important insights for further development. For example, to improve accessibility, particularly in resource-limited settings, we identified a need to reduce costs further (from approx. 700 €) and streamline the user experience by moving from disjointed software tools to a cohesive, AI-integrated workflow.

This paper introduces the Entomoscope 2.0, an open-source platform that automates the workflow of insect digitization, from image capture to analysis and data sharing. The platform combines affordable (400 €), user-friendly hardware with a new software suite, ENIMAS 2.0, offering enhanced image processing and morphometric tools. Its main advances include: (1) a validated, low-cost design that simplifies assembly and routine use; (2) an AI-based workflow that automates specimen imaging, curation, morphometric measurements, and basic taxonomic classification; (3) a case study demonstrating significant speed advantages over commercial workflows; and (4) a modular plugin system that supports high-throughput batch processing, scalable parallelization and community-driven extensions, ensuring the user community does not depend on a single research group but can integrate new code and algorithms independently.

## Methods

### Hardware Design and Fabrication

The main reasons for developing the Entomoscope 2.0 were cost reduction, standardization, further simplification, and additional software features. We thus used only ISO (International Organization for Standardization) or DIN (*Deutsches Institut für Normung*) standardized parts to profit from the market, reducing the prices of components available from different suppliers. As a side effect, this made sourcing parts from all over the world easier. In addition, we aim to reduce the number of required parts and components, first for off-the-shelf components, and also for 3D-printed components. The main functionality should stay the same, to make both systems interchangeable and comparable. As described for the Plug-In Entomoscope (Wührl et al., 2024), it should feature an automated linear stage to move the camera up and down for focus stacking and to make the system adaptable for different-sized lenses with fixed focal length and working distance, and therefore, no focus or zoom. In addition, we received a lot of valuable feedback on the Plug-In Entomoscope, which is in use at 13 entomology labs in six countries and has been used to digitize >10,000 specimens. This feedback was incorporated as design requirements. The two most obvious changes concern the lighting area. First, enlarging the lighting area allows moving the Petri dish rather than the specimen itself. This makes it easier to position the specimen in the field of view, especially for small specimens. Second, lowering the lighting area to table height allows for a more comfortable working position when taking many images. The resulting hardware platform is shown in Fig. 1.

**Fig. 1.**
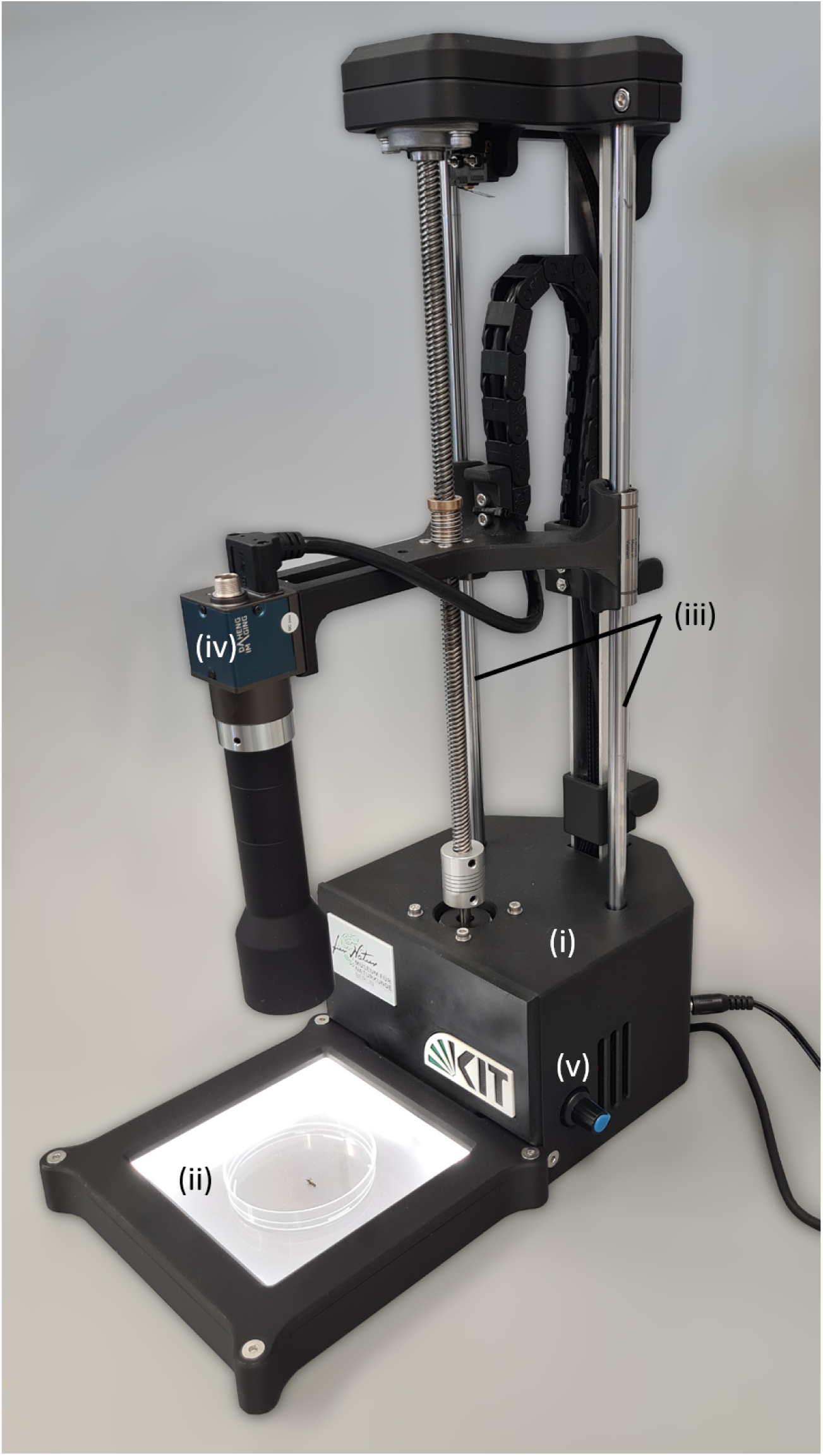
The Entomoscope 2.0 with (i) base, (ii) lighting, (iii) linear stage, (iv) USB camera, (v) dimmer knob.

### Software Platform: ENIMAS 2.0

The second-generation Entomoscope is powered by an AI-integrated software suite, ENIMAS 2.0 (ENtomoscope IMAging Software v2.0). This new version marks a fundamental paradigm shift from the original hardware control utility to a comprehensive, AI-powered analytical platform. The software is developed in Python 3 and features a graphical user interface (GUI) (Fig. 2) built with the PyQt5 framework, ensuring an intuitive user experience for researchers without programming expertise. The architecture is designed to support a seamless end-to-end workflow, from specimen imaging to data analysis and sharing.

**Fig. 2.**
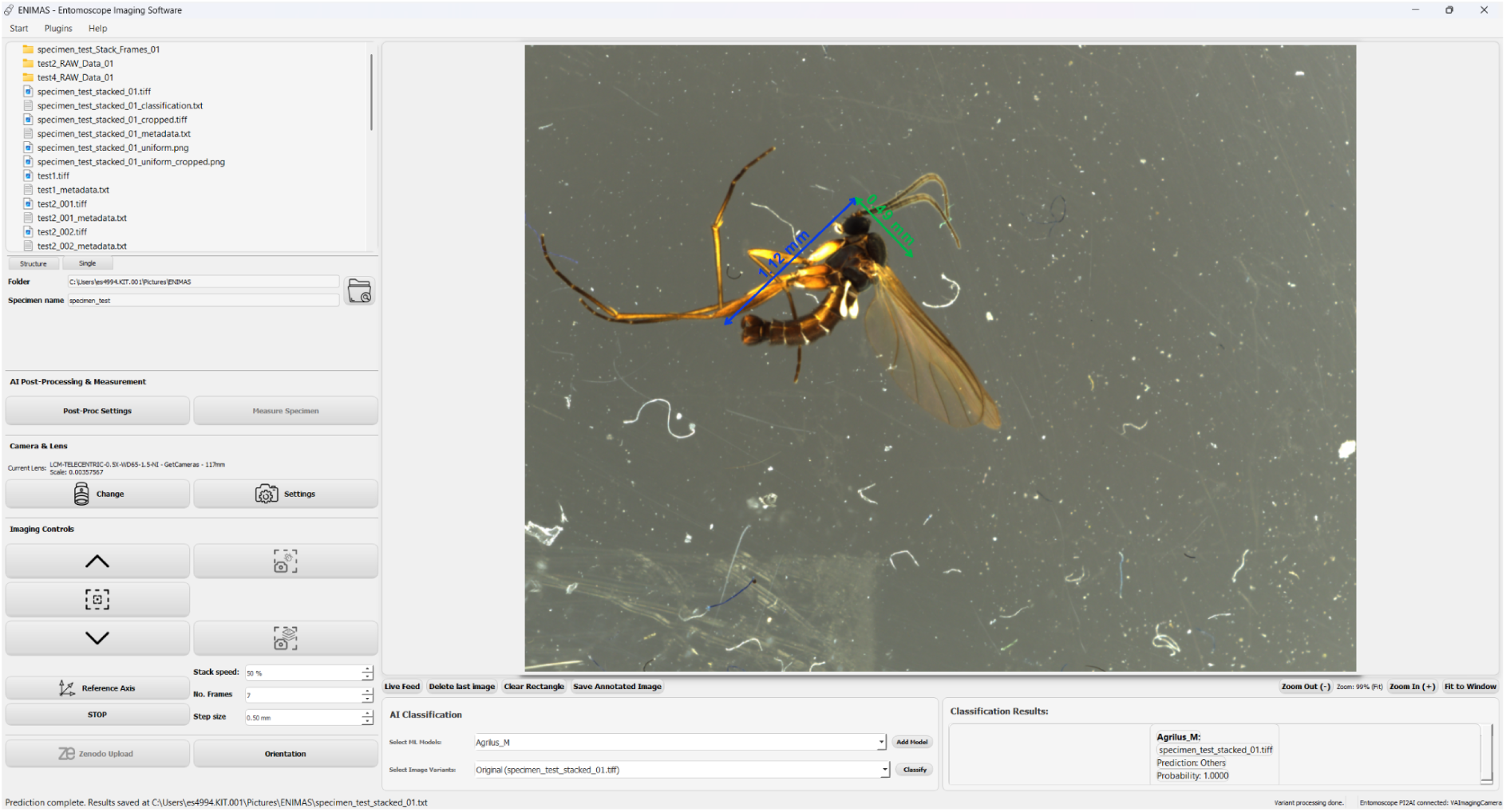
The ENIMAS 2.0 Graphical User Interface (GUI). The screenshot shows the integrated AI workflow, with an automated Oriented Bounding Box (OBB) measurement (length and width) applied to a specimen.

### Installation and Accessibility

To ensure the platform is accessible to its target audience of researchers and citizen scientists, who may not have programming expertise, we developed a fully automated installation script for Windows (10/11). This single batch file (install.bat) handles the complete setup process: it installs the correct Python version (3.10.8), sets up an isolated virtual environment, and installs all required dependencies, including PyTorch (CPU-only build) and OpenCV. Critically, the script also automatically detects and installs the necessary hardware drivers for both ArduCam and VAImaging camera systems. Upon completion, a desktop shortcut is created that launches the application, bypassing any need for command-line interaction. This streamlined, one-click process, supplemented by a comprehensive User_Guide.pdf (see Code and Data Availability), allows a user to turn a standard computer into a fully functional imaging station in approximately 10-15 minutes.

### Core Functionality and Image Acquisition

To provide acquisition flexibility, the software offers two image acquisition modes. For relatively flat or rapidly digitizable specimens, the user can acquire a single high-resolution image; for larger specimens that require greater depth of field, the software performs automated focus stacking. In this mode, ENIMAS 2.0 interfaces directly with the linear stage to capture a series of images at precise vertical intervals. To generate the final composite image from this stack, the user can select one of three processing methods:

1. **Helicon Focus Integration:** The software can automate the commercial program Helicon Focus (Helicon Soft, 2025) by running it automatically in the background. This method provides industry-standard, high-quality stacking results.
2. **Built-in Stacking:** A fully self-contained, open-source algorithm is provided. This method first aligns images using the Enhanced Correlation Coefficient (ECC) technique (Evangelidis and Psarakis, 2008). Then it fuses the stack using a Laplacian-based operator to select the sharpest pixel regions in each frame.
3. **No Stacking:** This mode saves all individual images from the stack without processing, allowing for manual post-processing or quality control.

The output from any of these paths (a single shot, a composite image, or a stack of images) serves as input to the subsequent analysis pipeline.

### AI-Powered Analysis Pipeline

A key innovation in ENIMAS 2.0 is the integration of a multi-stage AI pipeline that automates post-acquisition processing.

- **Automated Specimen Cropping:** To isolate the specimen from the unprocessed image, ENIMAS 2.0 implements a hybrid workflow that allows the user to select between two methods based on their requirements for speed or precision.
- The first method, “YOLO-Fast”, utilizes an object detection model based on the YOLOv8 architecture (Jocher et al., 2023). We fine-tuned this model on a custom dataset of 257 insect images, which were manually labeled for this study. To facilitate the training of future models, this annotated dataset is made publicly available in the supplementary materials (Zenodo).
- The second method, “BoxSegmenter-Accurate”, is a training-free approach designed for high-fidelity cropping. This method employs the BoxSegmenter model from the Finegrain AI Refiners library (finegrain-ai/refiners, 2025). Instead of relying on class-specific training, it performs semantic segmentation to generate a precise pixel-level mask of the foreground object. The system then identifies the outermost contour of this mask and computes a tight, axis-aligned bounding box, ensuring accurate cropping even for specimens with complex or irregular boundaries.
- **Uniform Background:** For further standardization, the platform leverages the same BoxSegmenter segmentation model used in the ‘BoxSegmenter-Accurate’ method. The model generates a precise binary mask to isolate the specimen, which is then used as an alpha channel (transparency layer). This allows the original background to be algorithmically replaced with a user-selected uniform color (e.g., neutral gray, white, black, or 8 other presets) or a custom user-provided image. This process is critical for creating consistent datasets for machine learning applications and standardizing images for publication.
- **Automated Morphometrics with Human-in-the-Loop Supervision:** The system implements an automated measurement workflow based on the Oriented Bounding Box (OBB) method. This method uses a separate YOLOv8-obb model, specifically fine-tuned for oriented object detection, to detect the specimen, leveraging an architecture previously validated by Shirali et al. (2025). In this study, the model demonstrated high precision relative to manual Zeiss microscope measurements, achieving a Mean Absolute Error (MAE) of 0.2 mm and a Pearson correlation coefficient of R = 0.99 (approximately 2.3% relative error). Unlike a standard axis-aligned bounding box, the OBB can rotate to enclose the specimen at any orientation tightly. This is crucial for accurately deriving morphometric data, as the system can estimate the actual length and width from the box’s primary axes, regardless of the insect’s orientation. To ensure accuracy and user control, ENIMAS 2.0 integrates this automated method into a human-in-the-loop (HITL) workflow. The GUI allows the user to inspect the automatically generated OBB visually and, if necessary, manually draw a new box from scratch or adjust the existing box’s position, angle, and dimensions.
- **Real-Time Classification:** ENIMAS 2.0 features a flexible inference engine that leverages the ONNX (Open Neural Network Exchange) runtime. This design supports models trained in major deep learning frameworks, including PyTorch, TensorFlow, and Ultralytics YOLO, once they are converted to the standard ONNX format. The system is designed for high flexibility, allowing users to add their own pre-trained models via a simple GUI. When a classification is run, the top-1 predicted class and confidence score are displayed directly in the GUI, along with a convenient link to the expected class’s Wikipedia page (if available) for immediate information on the species. The engine also supports ensemble predictions, in which the outputs of multiple models are averaged to improve accuracy. Once a model is loaded, the software performs real-time inference on the captured image, displaying the top-1 predicted class and confidence score. To fully support reproducibility and extension by other researchers, a comprehensive technical guide (CLASSIFICATION_MODEL_GUIDE.pdf) is provided, detailing the complete workflow for training, formatting (including embedding metadata in the ONNX file), and integrating new custom models into the software.

### Modular Add-on Functionality

To maximize flexibility and provide extensible add-on functionality, we implemented a modular plugin architecture. This design decouples the core analysis functions from the Entomoscope’s image capture workflow, enabling high-throughput analysis of large, pre-existing image collections. Users can activate plugins to apply the same AI tools in batches to entire folders of images. To demonstrate this capability, we developed four initial plugins, one for each core AI function: automated cropping, uniform background generation, measurement, and classification. This architecture allows for the easy addition of new community-developed or user-specific analysis tools in the future without altering the core application.

### Data Management and Open Science Integration

The platform’s data management capabilities have been significantly expanded to support the new analytical workflows. For each specimen, ENIMAS 2.0 allows the user to save a comprehensive set of outputs to document the entire process. This includes the unprocessed image stack (if captured), the final composite image, various processed versions (e.g., cropped, with uniform background, or both combined), and an annotated image showing the measurement overlay. Crucially, it also generates a structured text file (.txt) containing all associated metadata, including OBB coordinates, length and width measurements, camera settings, and any classification results. To promote FAIR (Findable, Accessible, Interoperable, and Reusable) data principles and open science practices (Wilkinson et al., 2016), the software includes a one-click “Zenodo Upload” feature. This allows users to easily archive their complete datasets to Zenodo, a general-purpose open-access repository operated by CERN, which assigns a persistent Digital Object Identifier (DOI) to the dataset, making it formally citable.

### Validation Methodology

To evaluate the workflow efficiency of the Entomoscope 2.0, we conducted a comparative case study against a high-end commercial imaging system (Keyence VHX-7000/7100 series). It is important to note that the objective of this comparison was not to benchmark absolute optical superiority but to evaluate the operational scalability of the digitization workflow. Specifically, we aimed to demonstrate how low-cost, automated stations can enable high-throughput digitization by allowing laboratories to deploy multiple units in parallel (e.g., employing 5-10 students simultaneously for bulk sample sorting), a strategy that is cost-prohibitive with expensive commercial systems.

The study used 54 insect specimens randomly sampled from a bulk malaise trap catch. This sample size was determined to be sufficient to establish statistical significance, given the anticipated large effect size between the manual and automated workflows. To ensure the evaluation reflected real-world conditions for biodiversity monitoring, specimens were selected without prior taxonomic screening. The sample set represents a morphologically diverse collection, ranging in body length from approximately 1 to 4 mm. The workflow for each system was timed from initial imaging to the final processed output, including measurement, cropping, and background removal, as illustrated in Fig. 3.

**Fig. 3.**
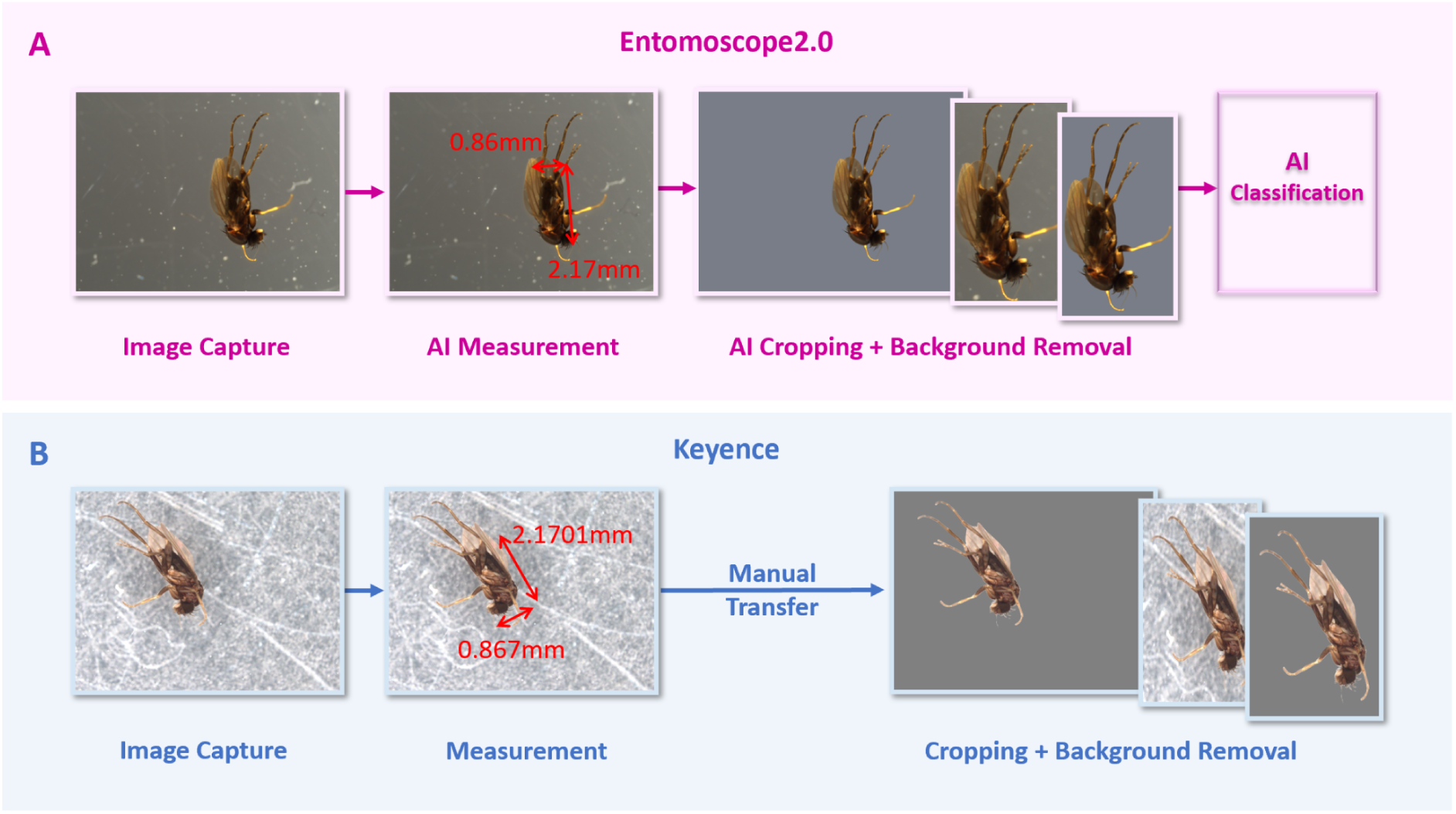
Digitization workflow comparison. (A) The single-step, integrated AI pipeline of Entomoscope 2.0. (B) The multi-step workflow of the Keyence system requires manual measurement and separate post-processing.

### Entomoscope 2.0: AI-Integrated Workflow

A single image was captured for each specimen. Immediately following acquisition, all analysis tasks, AI-powered measurement (OBB), cropping, and background removal were performed as a single, continuous process within ENIMAS 2.0. For this comparison, we used the “BoxSegmenter-Accurate” method for cropping and background removal to ensure high-fidelity output. Classification was excluded from this specific comparison to maintain parity with the Keyence system, which lacks this feature. The total time for this integrated workflow was recorded. All processing was performed on a laptop with a 13th Gen Intel Core i7-1370P CPU.

### Keyence: Standard Commercial Workflow

To represent the operational reality for biodiversity researchers, we utilized the standard workflow imposed by the Keyence system. Because the commercial software is a closed, general-purpose ecosystem, it does not natively support domain-specific tasks such as biological semantic segmentation or automated specimen cropping. Consequently, this workflow is inherently multi-stage:

- Step 1 (Acquisition & Manual Measurement): A single image was captured, and the user manually measured length and width using the system’s built-in tools (two separate actions). The time for this step was recorded.
- Step 2 (External Post-Processing): To achieve the same output as the Entomoscope, the captured images were transferred and processed on the same laptop using Windows 11 Photo Editor’s AI features for cropping and background removal. This tool was selected because its AI-driven segmentation is robust to background artifacts (e.g., scratches, dust). In contrast, standard thresholding tools (e.g., ImageJ) would require extensive custom scripting to achieve comparable results.

The total processing time for the Keyence workflow was the sum of Step 1 and Step 2. The time required for manual data transfer between systems was not included, making this a conservative time comparison.

### AI Plugin Batch Processing

Additionally, to quantify the performance of the ENIMAS 2.0 AI tools in a high-throughput context, the 54 captured images were processed in batch mode using the individual AI plugins: Cropping (YOLO-Fast), Cropping (BoxSegmenter-Accurate), Background Removal, Measurement (OBB), and Classification (YOLO11x). The total time for each discrete batch operation was recorded. All output images are provided in the supplementary materials for manual review.

### Statistical Analysis

To validate workflow efficiency, we compared processing times using statistical methods. First, we assessed the normality of the data distributions. Due to the non-normal distribution of the Entomoscope dataset, we utilized the Wilcoxon signed-rank test (paired). Unlike a standard t-test, this nonparametric test assesses whether the processing time was consistently lower for the Entomoscope across paired specimen measurements. To understand the magnitude of the improvement, we calculated Cohen’s d (effect size). While a p-value indicates if a difference exists, Cohen’s d quantifies how large that difference is relative to the variance (with d > 0.8 typically considered a “large” effect). Finally, to account for variability and provide a robust estimate of the expected performance gain in future applications, we computed the 95% Confidence Interval (CI) for the speedup factor using a bootstrapping method (n=10,000 resamples).

## Results

### The Integrated Entomoscope 2.0 Platform

The redesigned hardware and ENIMAS 2.0 software form a cohesive, user-friendly platform for insect digitization. The physical system provides a stable and ergonomic station for specimen imaging, while the software’s graphical user interface (GUI) provides centralized control over the entire integrated workflow, from image acquisition to final data export.

### Hardware Performance and Cost Reduction

The hardware architecture underwent a comprehensive redesign, reducing the total component cost to 400 €. This significant cost reduction was primarily achieved by strategically replacing custom or specialized components with readily available ISO- and DIN-standardized parts. Furthermore, the design was meticulously optimized for additive manufacturing (3D printing), reducing the number of distinct printed parts from 17 in the original Plug-In Entomoscope to 12 in Entomoscope 2.0, while keeping printing time similar (14 hours 25 minutes vs. 14 hours 49 minutes). Additional modifications included eliminating an extra USB-to-USB connector, with the internal USB hub’s cable now routed directly to the exterior. The system’s overall size has also reduced, with the base being 140 x 140 x 95 mm (Fig. 1 - i). The lighting area (ii) is now positioned at desk level and offers, therefore, enhanced ergonomic comfort. For the Entomoscope 2.0, threaded inserts were substituted with ISO-standardized hex nuts, and the previously used linear rail was replaced by linear rods integrated with linear ball bearings (iii). In response to component availability and to leverage advancements in imaging technology, an alternative camera module (MER2-1220-32U3C, manufactured by VAImaging) was incorporated (iv). This camera features an IMX226 sensor, which maintains the same high 12-megapixel resolution as the Raspberry Pi HQ Camera (IMX477 sensor) used in the Plug-In Entomoscope, while offering slightly larger individual pixels. A critical enhancement of the newly integrated camera is its native USB 3.0 connectivity, which eliminates the need for an additional USB-Board. This direct integration significantly improves system cohesion and operational simplicity, contributing to a more robust and efficient imaging platform. The resulting image quality, as produced by the IMX226 sensor, is comparable (e.g., in resolution) to or surpasses (particularly in terms of low-light performance, signal-to-noise ratio, and overall sensitivity) that of the original system, thereby ensuring the acquisition of high-resolution images suitable for rigorous morphological analyses and classification.

The Entomoscope 2.0 still allows varying the light intensity with a dimmer knob (v) installed on the right side of the base. The previously used custom dimmer board was replaced by an off-the-shelf board to reduce the system’s setup time. In general, the new design’s setup is much simpler than that of the previous version. Depending on the person in charge’s skill, we reduced the time to set up the system by approximately 50%. The system can now be set up in around 6-8 hours, if all parts are printed and available.

### Case Study: Workflow Efficiency Comparison

#### Integrated Workflow vs. Multi-Step Commercial System

The comparative time study revealed significant efficiency gains with the Entomoscope 2.0 (Table 1). The 54 specimens were processed from image capture to data acquisition in 2950.09 seconds (49.17 minutes), while the Keyence system, including manual post-processing, required 6713.42 seconds (111.89 minutes). Statistical analysis confirmed that the efficiency gain is highly significant (Wilcoxon signed-rank test, W = 0, p < 0.001). The study yielded a Cohen’s d of 2.98, indicating an extremely large effect size. Notably, the Entomoscope 2.0 workflow was faster than the Keyence workflow for all specimens tested (N=54). A complete statistical breakdown is provided in Table 1.

**Table 1.**
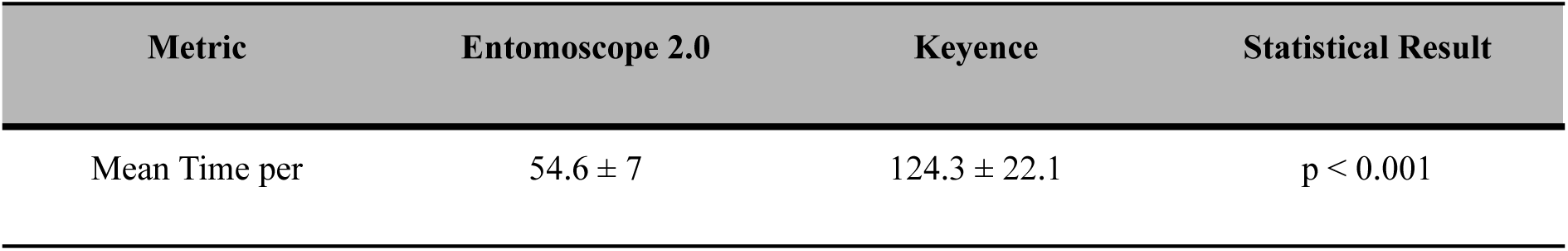
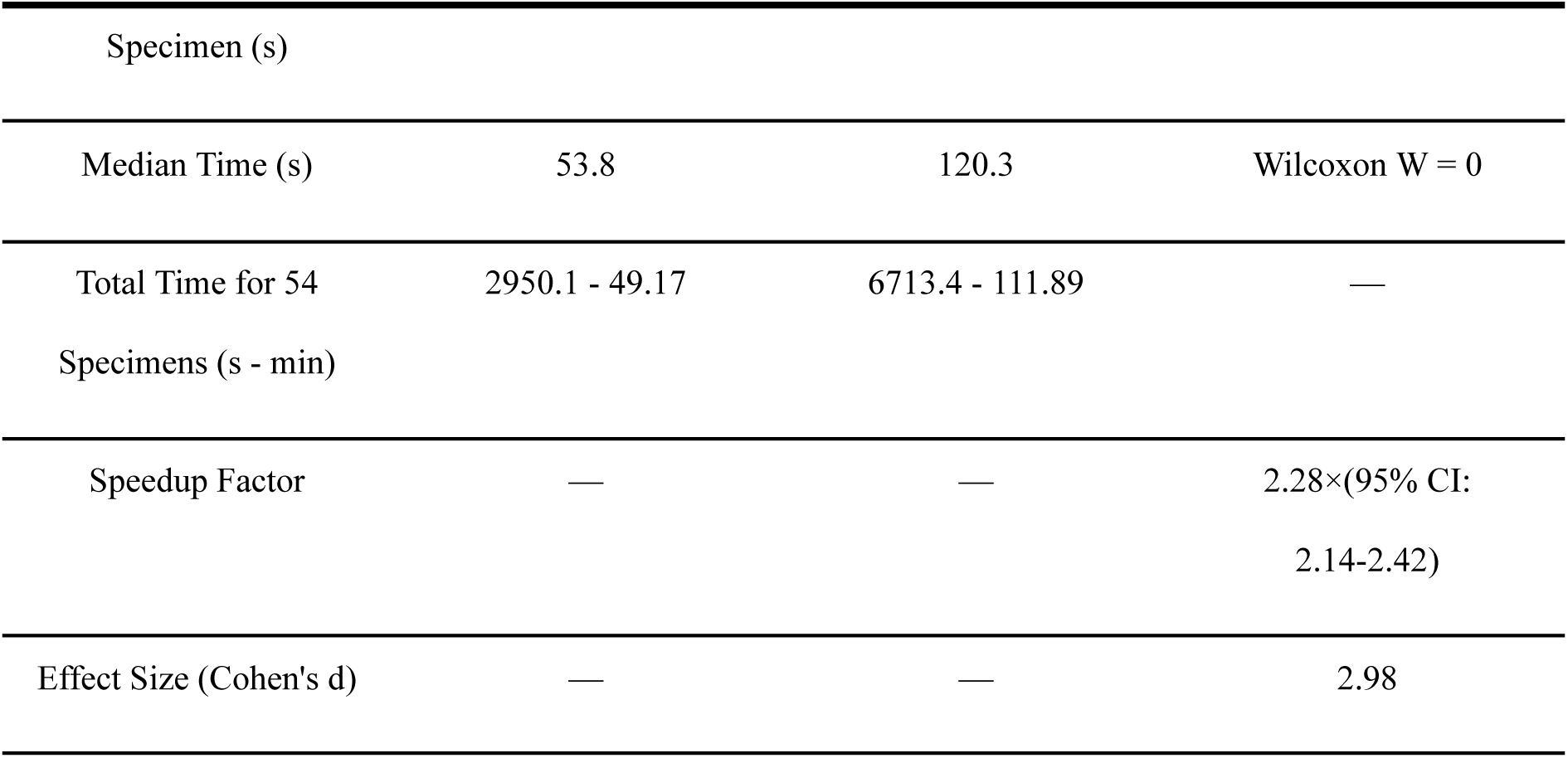
Statistical comparison of workflows (N=54). Key performance metrics for Entomoscope 2.0 vs. Keyence, showing a 2.28× speedup factor and high statistical significance.

| Metric | Entomoscope 2.0 | Keyence | Statistical Result |
| --- | --- | --- | --- |
| Mean Time per | $54.6 \pm 7$ | $124.3 \pm 22.1$ | $p < 0.001$ |
| Specimen (s) |  |  |  |
| Median Time (s) | 53.8 | 120.3 | Wilcoxon W = 0 |
| Total Time for 54 Specimens (s - min) | 2950.1 - 49.17 | 6713.4 - 111.89 | — |
| Speedup Factor | — | — | 2.28×(95% CI:<br>2.14-2.42) |
| Effect Size (Cohen's d) | — | — | 2.98 |

The mean processing time per specimen was 54.6 seconds for Entomoscope 2.0, compared to 124.3 seconds for the Keyence workflow (Fig. 4C). The per-specimen comparison (Fig. 4A) indicates that the Entomoscope 2.0 was faster in all cases. To quantify this, we analyzed paired measurements (Fig. 4B), which revealed a universally positive slope for all specimens (N=54). The distribution analysis (Fig. 4C) further highlights the performance stability: the Entomoscope 2.0 workflow exhibited a tight variance around the median (M=53.88 s) with a standard deviation of only 7 s. This low variance reflects the high reliability of the automated OBB model, as manual interventions (HITL) were rarely needed during validation. In contrast, the Keyence workflow displayed a broad, right-skewed distribution (M = 120.3 s, Std Dev = 22.1 s) with significant outliers attributable to interventions during manual measurement or post-processing.

**Fig. 4.**
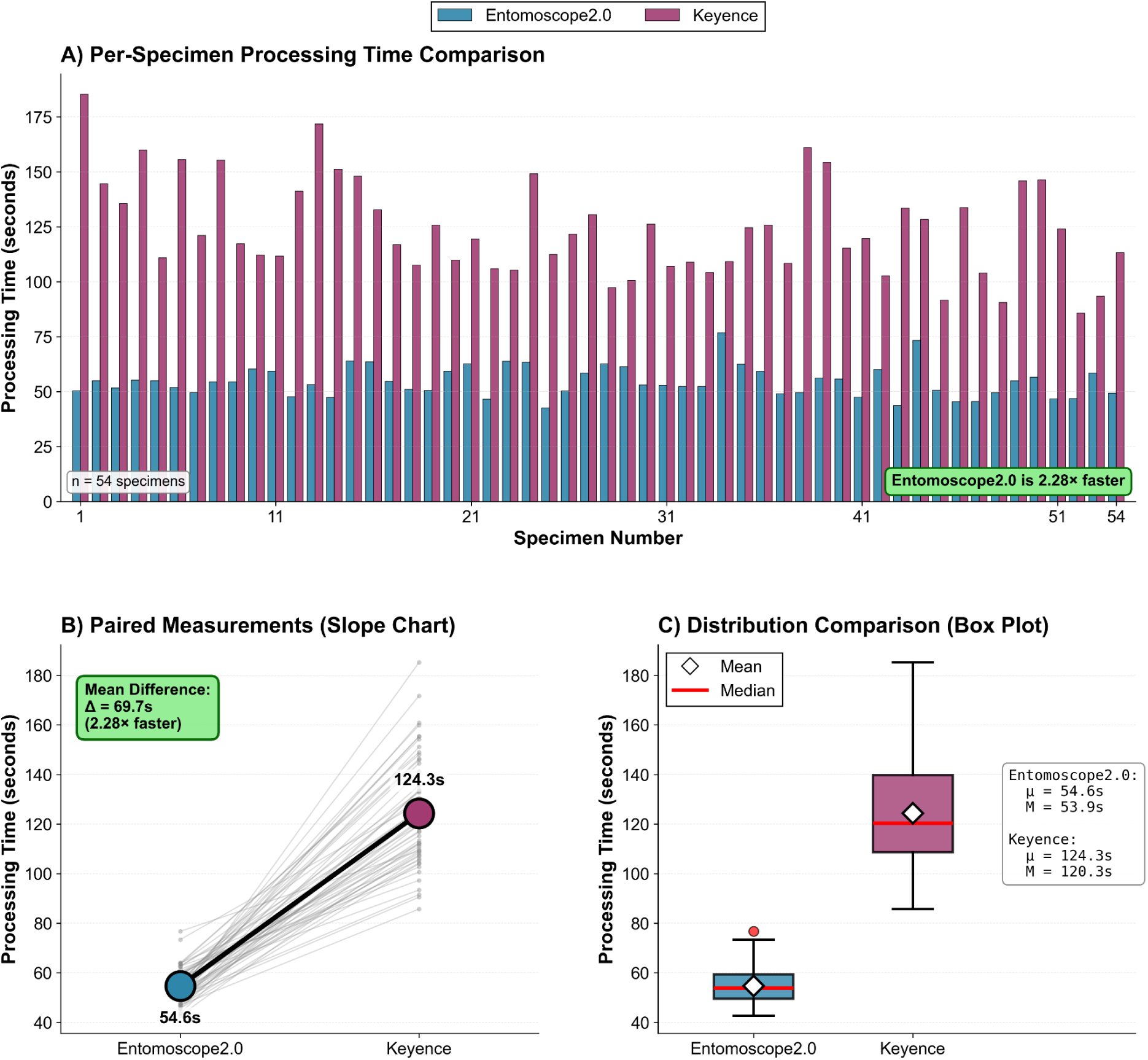
Comparative workflow efficiency (N=54). (A) Per-specimen processing time for Entomoscope 2.0 (blue) vs. Keyence (purple), showing consistently faster performance for the Entomoscope across all samples. (B) Paired slope chart illustrating the time difference for each specimen; the positive slope indicates the Entomoscope was faster in every instance. (C) Box plot comparison of time distributions. The Entomoscope 2.0 workflow (Mean = 54.6 s, SD = 7.0 s) is significantly faster and exhibits lower variance (SD = 7.0 s) than the Keyence workflow (Mean = 124.3 s, SD = 22.1 s), resulting in a 2.28× speedup.

### AI Plugin Batch Processing Performance

To assess the platform’s utility for high-throughput analysis of existing image collections, all specimen images were processed using the standalone AI plugins in batch mode. The results (Fig. 5) highlight the performance characteristics of each AI tool.

**Fig. 5.**
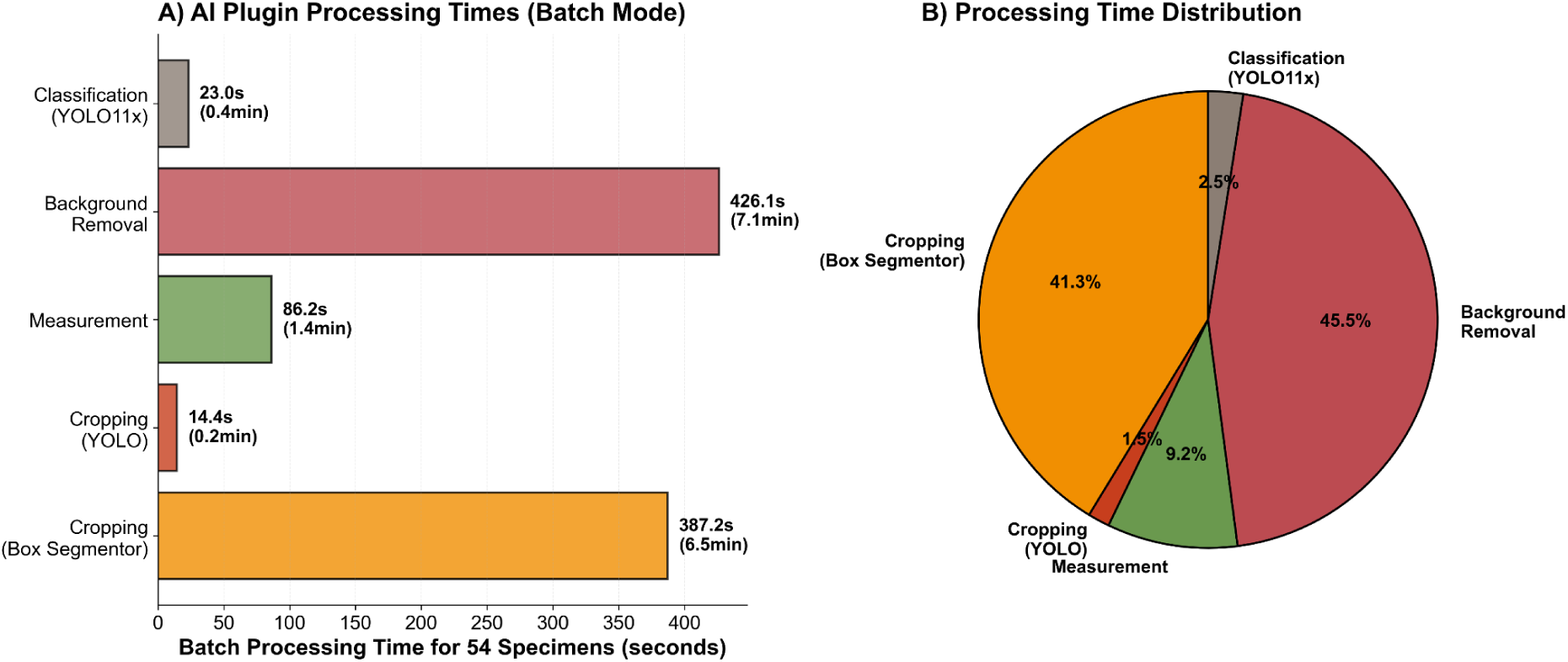
AI plugin batch processing performance (N=54). (A) Total processing time (seconds) (B) Proportional time distribution for each AI module.

The “YOLO-Fast” cropping plugin was exceptionally fast, processing the entire batch in just 14.4 seconds (0.2 min), demonstrating its suitability for large-scale, rapid processing. The more precise “BoxSegmenter-Accurate” cropping method took 387.2 seconds (6.5 min). The most computationally intensive task was background removal, requiring 426.1 seconds (7.1 min). The AI-powered (OBB) measurement plugin processed the 54 images in 86.2 seconds (1.4 min). The classification plugin, demonstrated using a large general-purpose model (YOLO11x) to benchmark inference efficiency, completed the batch in 23.0 seconds (0.4 min). We emphasize that this metric reflects computational throughput (inference speed) rather than predictive accuracy, which is dependent on the specific user-provided model. This processing time would be even shorter with lighter model architectures.

### Extrapolated Efficiency at Scale

Fig. 6 illustrates the baseline time savings when comparing a single Entomoscope 2.0 unit against a single commercial system. Even in this 1-to-1 comparison, the efficiency gains are substantial: for a project of 10,000 specimens, a single Entomoscope saves over 190 hours compared to the commercial workflow.

**Fig. 6.**
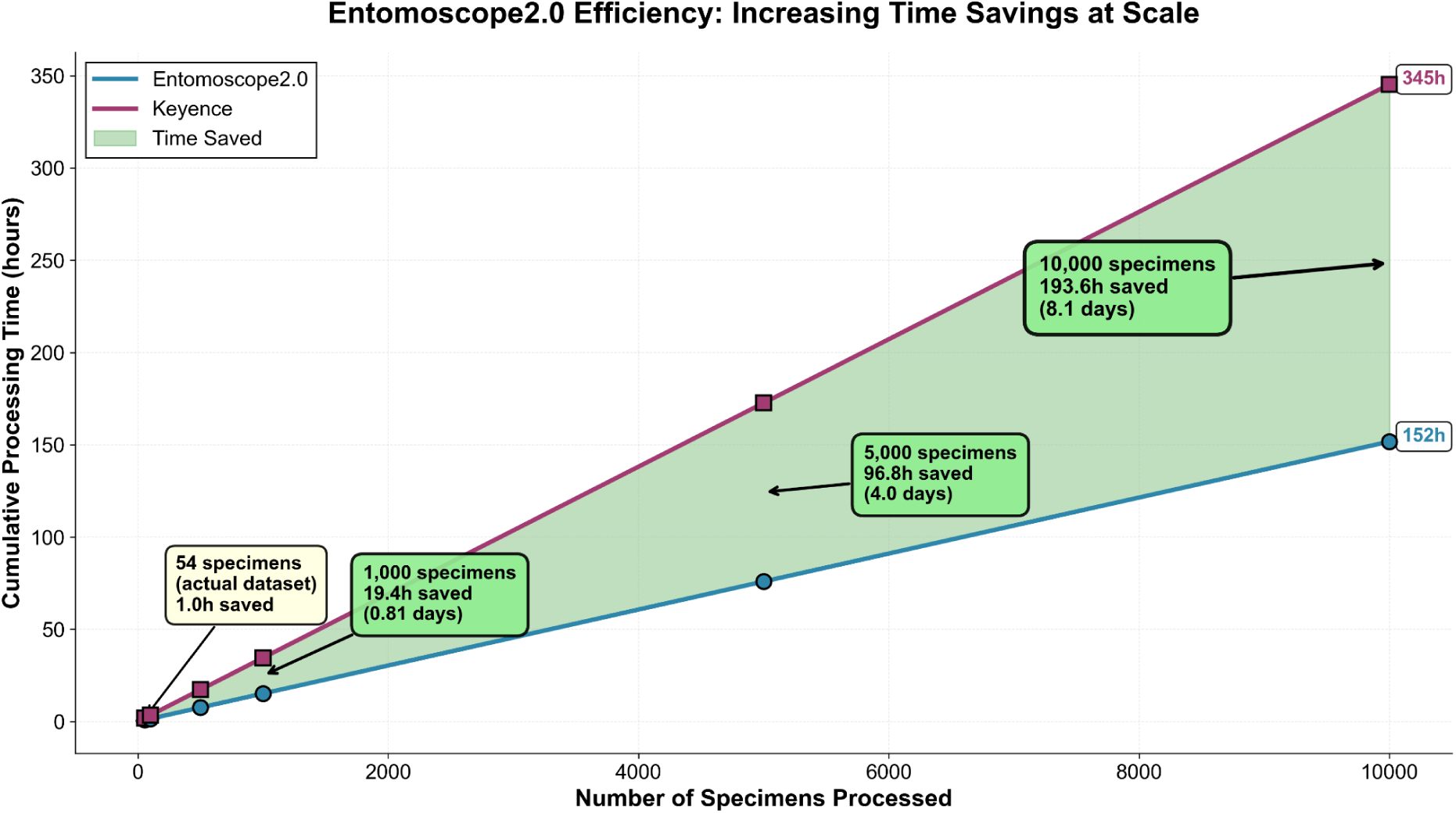
Extrapolated time savings at scale. The shaded area shows cumulative time saved by Entomoscope 2.0. At 10,000 specimens, the system saves a projected 193.6 hours (8.1 days) of processing.

However, this linear comparison understates the platform’s true scalability. Commercial systems like the Keyence VHX-7000 represent a massive capital investment (typically €50,000–€80,000), effectively limiting most laboratories to a single unit. This creates a problematic bottleneck: specimens must be processed serially, one by one.

In contrast, the low material cost of the Entomoscope 2.0 (400 €) fundamentally alters the economics of digitization. For less than 5% of the cost of a single commercial system, a laboratory could build 10 Entomoscope units. This allows a parallelized workflow in which multiple operators—such as a team of students or parataxonomists—can digitize specimens simultaneously.

If we apply this parallelization to the scenario in Fig. 6, deploying just five Entomoscope units would divide the total project timeline by a factor of five. A digitization task that would take 345 hours (approx. 8.5 weeks) on a single commercial system could be completed in just 30 hours (< 1 week) using a small array of Entomoscopes. This massive scalability positions the Entomoscope 2.0 as a uniquely capable solution for tackling the immense backlog of “dark taxa” and bulk monitoring samples.

## Discussion

The accelerating biodiversity crisis has generated an urgent need to characterize millions of insect specimens collected in bulk monitoring campaigns. Processing these samples requires a fundamental shift from artisanal, single-specimen imaging to industrial-scale digitization. The primary contribution of the Entomoscope 2.0 is not merely cost reduction, but the enablement of a new paradigm: parallelized specimen processing at scale.

While high-end commercial systems offer exceptional optical quality and versatility, they are often proprietary, cost-prohibitive, and characterized by a large physical footprint. As laboratory space is often as limited as funding, relying on large, centralized workstations creates a bottleneck where thousands of specimens must wait for serial processing on a single machine. By contrast, the Entomoscope 2.0’s compact design and low cost (400€) allow laboratories to maximize their limited bench space. A standard laboratory bench that holds one large commercial microscope can easily accommodate 5–10 Entomoscope units. This allows a small team to process bulk samples in parallel, effectively multiplying throughput without requiring “endlessly large” laboratory facilities. This scalability is essential for generating the massive, standardized image datasets needed to train the next generation of AI identification models.

While several low-cost open-source microscopy solutions have been proposed, they primarily focus on hardware accessibility and manufacturing innovation. For instance, Salido et al. (Salido et al., 2022) reviewed the landscape of open hardware microscopy, highlighting the trade-off between portability and automation, while Christopher et al. (Christopher et al., 2025b) recently demonstrated the feasibility of a fully 3D-printed optical system, including lenses. The Entomoscope 2.0 complements these hardware advancements by integrating a low-cost imaging station with a specialized, domain-specific AI workflow. Unlike systems that focus solely on image acquisition, our platform provides an end-to-end solution—from capture to a curated dataset—uniquely positioning it to bridge the gap between affordable hardware and actionable biological data.

Furthermore, the case study results, comparing our system to a high-end Keyence platform, show a significant 2.28-fold increase in workflow efficiency. This acceleration is achieved by replacing a multi-step process—involving manual data transfer and separate post-processing tools—with a single, cohesive, AI-powered workflow (Fig. 3). It is also important to note that this 2.28-fold speedup was achieved using a standard laptop CPU (13th Gen Intel Core i7-1370P, 1.90 GHz) for all computations. This represents a baseline, as ENIMAS 2.0 is designed to be accessible without requiring a dedicated GPU. Performance could be further accelerated on systems where optional GPU support for the AI models is enabled. Crucially, this comparison is not a critique of commercial hardware, which is optimized for extreme resolution and broad applicability. Instead, it highlights the trade-off between a general-purpose manual workflow and a specialized automated pipeline. Commercial systems, designed to handle everything from material science to biological imaging, often rely on manual measurement tools and external post-processing software that are not specific to entomology.

The efficiency gain is not only quantitative but qualitative. The commercial workflow required manual morphometric measurements, which involved two distinct user actions (drawing a line for length, then another for width). In contrast, Entomoscope 2.0’s AI-based OBB method captures both metrics with a single automated detection, shifting the user’s role from manual labor to simple verification (HITL). While the computational accuracy of the underlying OBB algorithm is established (MAE = 0.2 mm (Shirali et al., 2025)), the primary advantage in the Entomoscope 2.0 workflow is the integration of this precision with Human-in-the-Loop (HITL) supervision. By presenting the user with a pre-calculated, high-confidence bounding box, the system shifts the user’s role from manual drawing to rapid verification. This ensures that the final measurement accuracy remains equivalent to manual expert ground truth, as the user corrects any algorithmic deviations, while retaining the significant speed advantage of automation.

Moreover, the reliance on third-party AI tools (e.g., Windows 11 Photo Editor) for post-processing in the Keyence workflow introduced performance inconsistencies. We qualitatively observed that these general-purpose tools, while effective for standard photography, occasionally struggled with complex insect morphology (e.g., appendages), requiring time-consuming manual correction. This aligns with the high variance (Std Dev = 22.1 s) and time spikes seen in the Keyence data. ENIMAS 2.0’s dedicated models (BoxSegmenter and YOLO), trained and optimized for insect digitization, deliver consistent, reliable performance, as evidenced by the low variance (Std Dev = 7 s) in the Entomoscope 2.0 workflow.

The batch processing results (Fig. 5) also provide a clear guide for users: the “YOLO-Fast” cropping plugin offers remarkable speed (14.4s for 54 images), making it ideal for large-scale data triage, while the “BoxSegmenter-Accurate” method provides a more precise, albeit slower, alternative for final curation. This modularity allows researchers to balance the trade-off between speed and precision for their specific needs.

While the Entomoscope 2.0 platform represents a significant step forward, we acknowledge its limitations, which provide clear avenues for future work. For example, the accuracy of rapid taxonomic screening depends entirely on the quality and scope of the user-provided classification model. The system’s reliance on the optional commercial software Helicon Focus for the highest-quality stacking introduces a minor cost barrier. However, this limitation is significantly mitigated by the inclusion of a built-in, open-source stacking algorithm (Section 2.2.2), enabling the platform to function fully without third-party software.

Future development will focus on addressing these points. A key priority is improving the performance of the native, open-source focus stacking algorithm to make it fully competitive with commercial options. We also plan to develop and train more robust, generalized classification models for common insect groups to provide robust “out-of-the-box” functionality. We will also finalize the one-click Upload feature that will allow for storing images in an open-access database. Finally, we aim to foster a user community around the platform, encouraging the development and sharing of new plugins and trained models through the modular architecture, thereby further enhancing the system’s capabilities and impact.

Beyond individual laboratories, we envision the Entomoscope 2.0 as a valuable tool for a distributed, global biodiversity monitoring network. By standardizing both the imaging hardware and metadata structure (in line with FAIR principles), the platform enables the creation of large-scale, interoperable datasets. This 1:1 compatibility supports global initiatives such as the proposed Global Repository of Insect Traits (GRIT) (Cardoso et al., 2025). Widespread adoption could enable the rapid digitization of ‘dark taxa,’ facilitate the detection of phenotypical shifts across geographic gradients, and generate the massive, standardized training datasets required to advance the next generation of AI-based taxonomic identification models.

## Conclusion

In summary, this paper presents the Entomoscope 2.0, a smart open-source platform that integrates optimized hardware for image acquisition with an AI-powered software suite for automated analysis. Our results demonstrate that the system achieves a 2.28-fold increase in workflow efficiency compared to a standard commercial pipeline. Crucially, the platform’s low cost (400€) and small footprint enable a new operational model: the deployment of parallelized imaging arrays to tackle the massive backlog of insect bulk samples. By democratizing access to high-throughput digitization, Entomoscope 2.0 empowers a global community of researchers, educators, and citizen scientists to contribute to the critical task of understanding, monitoring, and conserving insect biodiversity.

## Acknowledgments

Our work was supported by funding from the Center for Integrative Biodiversity Discovery at the Museum für Naturkunde Berlin and by grant #ZF4717901SK9 of the program Natural, Artificial and Cognitive Information Processing (NACIP) of the Helmholtz-Association, Germany.

During the preparation of this work, the authors used AI tools to improve clarity and fluency in the writing process. After using these tools, the authors reviewed and edited the content as needed and took full responsibility for the publication’s content.

## Data Availability

The ENIMAS 2.0 software is open-source and available on <u>GitLab</u>. The hardware design files, including CAD models, 3D-printing STL files, and the bill of materials, are available in the wiki section of the same GitLab repository. All validation data, including the test images and ground-truth measurements, are publicly archived on <u>Zenodo</u>. Comprehensive documentation, including detailed User_Guide.pdf and CLASSIFICATION_MODEL_GUIDE.pdf for training and integrating new classification models, is available in the GitLab repository.

